# *SlATG8f* enhances tomato thermotolerance and fruit quality via autophagy and HS pathways

**DOI:** 10.1101/2025.09.23.678159

**Authors:** Qunmei Cheng, Wen Xu, Cen Wen, Zhuo He, Liu Song

## Abstract

Plant responses to high-temperature stress involve dynamic and complex changes in physiology, biochemistry, and gene expression. Autophagy, a highly conserved intracellular degradation system, facilitates the removal and recycling of damaged cytoplasmic components such as proteins and organelles. It plays an essential role in plant growth, development, and adaptation to various stresses. Although the functions of many autophagy-related genes under high-temperature stress have been characterized, the role of *SlATG8f*, a key member of the *SlATG8* family in tomato, remains poorly understood. In this study, we generated *SlATG8f-*Overexpressing tomato lines using the recombinant vector pBWA(V)HS. Using quantitative reverse transcription polymerase chain reaction (qRT-PCR), we analyzed physiological indices and expression levels of *ATG8* family members and heat shock protein-related genes in fruits of wild-type (WT) and *SlATG8f*-overexpressing plants across four developmental stages under high-temperature stress. Our results indicate that overexpression of *SlATG8f* upregulates the expression of autophagy-related and heat shock protein genes, promotes early fruit ripening, and improves fruit quality under high-temperature stress. These findings reveal a regulatory role for *SlATG8f* in maintaining tomato fruit quality under heat stress and provide a theoretical foundation for tomato variety improvement.

## Introduction

Autophagy is an evolutionarily conserved process in eukaryotes that mediates the recycling of intracellular components and damaged macromolecules, playing a crucial role in plant development and stress adaptation[1]. Among the core autophagy-related (*ATG)* proteins, *ATG8* is particularly important for autophagosome formation and function. In plants, the *ATG8* gene family has expanded considerably from a single ancestral gene in algae to multiple isoforms in higher plants, reflecting its functional diversification[2]. Most autophagic receptors and adaptors contain an *ATG8*-interacting motif (AIM/UIM), which facilitates specific binding to *ATG8* proteins [3]. Growing evidence indicates that autophagy is activated under various abiotic stresses—such as nutrient deficiency, salinity, drought, and extreme temperatures—where it generally promotes cell survival and helps maintain cellular homeostasis[4]. However, the role of autophagy can vary depending on the type of stress, tissue, and developmental stage. For example, in barley, autophagy has been linked to programmed cell death in microspores under cold stress, and its inhibition resulted in reduced cell death [5].suggesting that autophagy can exert either pro-survival or pro-death functions depending on the context [6].

Heat stress (HS) is a major constraint on crop productivity and quality. It has been shown to induce the expression of autophagy-related genes and promote autophagosome formation [7]. The extent of autophagic activation depends on stress intensity, duration, and plant genotype. Studies using loss-of-function mutants of Arabidopsis (*ATG5* and *ATG7)* have revealed increased heat sensitivity, accompanied by more severe damage to photosynthetic and membrane systems[8]. Conversely, overexpression of certain ATG genes, such as *MdATG18a* in apple, enhanced thermotolerance by alleviating oxidative damage and protecting chloroplast function[9]. Several autophagy genes—including *ATG5, ATG6, ATG12a, ATG18a, ATG8a, ATG8c, ATG8g*, and *ATG8i*—have been implicated in the plant response to high-temperature stress [8,10,11]. However, the function of *SlATG8f*, a key member of the tomato *ATG8* family, under high-temperature conditions remains poorly understood.

Tomato (***Solanum lycopersicum L*.)** is a major greenhouse crop and an important model species for studying fruit development and stress responses. Heat stress (HS) severely impairs multiple reproductive processes, including pollen viability, fertilization, and fruit set, leading to significant yield reduction and quality losses [12,13]. Temperatures exceeding 35 °C disrupt key physiological and biochemical processes, adversely affecting sugar metabolism, pigment accumulation, and fruit firmness [14].Furthermore, HS alters root–shoot signaling and nutrient partitioning, further compromising fruit development [15].These detrimental effects underscore the need to elucidate the molecular mechanisms that could be targeted to improve thermotolerance.

In this study, we generated *SlATG8f-*overexpressing transgenic tomato lines and exposed them to continuous high-temperature stress (35 °C) during fruit development. Using quantitative reverse transcription PCR (qRT-PCR), we examined the expression profiles of *ATG8* family members and heat shock protein (HSP) genes in both wild-type and transgenic plants across four distinct fruit developmental stages. We also assessed key fruit quality parameters, including sensory properties (external color and firmness), flavor-associated traits (sugar-acid ratio), and health-promoting compounds (carotenoids). Our results indicate that *SlATG8f* overexpression enhances autophagic activity and upregulates HSP gene expression under heat stress, which promotes earlier fruit ripening and improves overall fruit quality. These findings offer novel insights into the function of *SlATG8f* in thermotolerance and fruit development, and provide a valuable genetic resource for breeding tomato varieties with enhanced heat resilience and superior fruit quality.

## Materials and methods

### Plant material handling and growing conditions

In this study,wild-type (WT) and *SlATG8f-OE* homozygous lines (F2 generation) of ‘Micro-Tom’ tomato were cultivated using a conventional soilless substrate. When the first truss of fruit reached the expansion stage, nine WT and nine transgenic plants were transferred to a climate-controlled growth chamber for heat treatment (35 °Cday/21 °Cnight), with plants grown at 25°Cday /16 °Cnight serving as the control[16], Fruit samples were collected at four developmental stages: mature green, breaker, orange ripe, and red ripe. Measurements included firmness, soluble solid content, titratable acidity, malondialdehyde (MDA) content, and the activities of key antioxidant enzymes. Additionally, the expression levels of *ATG8* family members and heat shock protein genes were analyzed by qRT-PCR. The number of seeds per fruit was recorded, and fruit color progression was monitored throughout ripening.

### Generation of tomato lines with *SlATG8f* overexpression

The *SlATG8f* overexpression vector was constructed by cloning the *SlATG8f* coding sequen ce into the pBWA(V)HS vector. Healthy tomato cotyledons were infected with Agrobacterium tumefaciens strain GV3101 harboring the 35S::*SlATG8f* construct. Transgenic lines were generated through Agrobacterium-mediated transformation. The expression level of *SlATG8f* in the overe xpression lines *(SlATG8f-OE)* was confirmed by qRT-PCR. Homozygous F2 plants were selected for subsequent experiments [17]

### Assessment of fruit color transition and fruit set rate

The timing of initial color change (recorded from day 0 of treatment) and the number of fruit set were documented through direct observation across different fruit samples.

### Analysis of fruit quality parameters and enzyme activities

Soluble solids content (SSC) and titratable acidity (TA) were measured using a PAL-BX/ACID1 refractometer (ATAGO, Japan). Carotenoid and malondialdehyde (MDA) levels, as well as activities of relevant enzymes and other physiological indicators, were quantified via UV spectrophotometry.Detailed experimental procedures were performed as described in previously establishedmethods:carotenoids [18], MDA[19],antioxidant enzyme activities [20].

### The expression levels of *SlATG8f* gene and HS gene were detected

The expression levels of *ATG8* family members and heat shock protein (HSP) response genes were quantified using real-time quantitative polymerase chain reaction (qRT-PCR). Total RNA was extracted from frozen tomato fruit samples with a Tengen RNA extraction kit (DP432, Tengen). Complementary DNA (cDNA) was synthesized using a Tengen reverse transcription kit (KR118), followed by qRT-PCR analysis. Gene-specific primers were designed with Premier 5.0 software and synthesized by Bio Company. The names and sequences of all primers are listed in Table S1. qRT-PCR was performed using the TB Green™ Premix Ex Taq™ II (Tli RNaseH Plus) kit (Takara) under the following conditions: each reaction was run in three technical replicates. The tomato Actin gene (*Solyc03g078400)* was used as the internal reference, and relative gene expression levels were calculated using the 2^-ΔΔCT^ method.

### Data processing and analysis

The experiment followed a completely randomized design with independent biological replicates. Each treatment group included three replicates, and random sampling was performed within each group. Data were organized using Microsoft Excel 2019 and analyzed with SPSS Statistics 23.0. Statistical significance was determined using Duncan’s multiple range test, with differences considered significant at P<0.05, indicated by different lowercase letters. Figures were generated using Origin 2022, and all data are presented as mean ± standard deviation.

## Results

### High temperature stress improves fruit thermotolerance in *SlATG8f-OE* plants

To evaluate the effect of *SlATG8f-* overexpression(*SlATG8f*-OE) on tomato fruit development under heat stress, fruit phenotypes were compared across treatment groups (Fig. 1). Under control conditions, no discernible differences were observed between wild-type (WT) and OE fruits. However, high-temperature stress induced fruit deformation in both genotypes, with more severe malformations evident in OE lines. Anatomical observations indicated that OE fruits accelerated and completed locular gel formation earlier than WT under normal conditions. Under heat stress, gel development was delayed in both WT and OE fruits, although OE plants exhibited greater tolerance compared to WT.

**Fig. 1.**
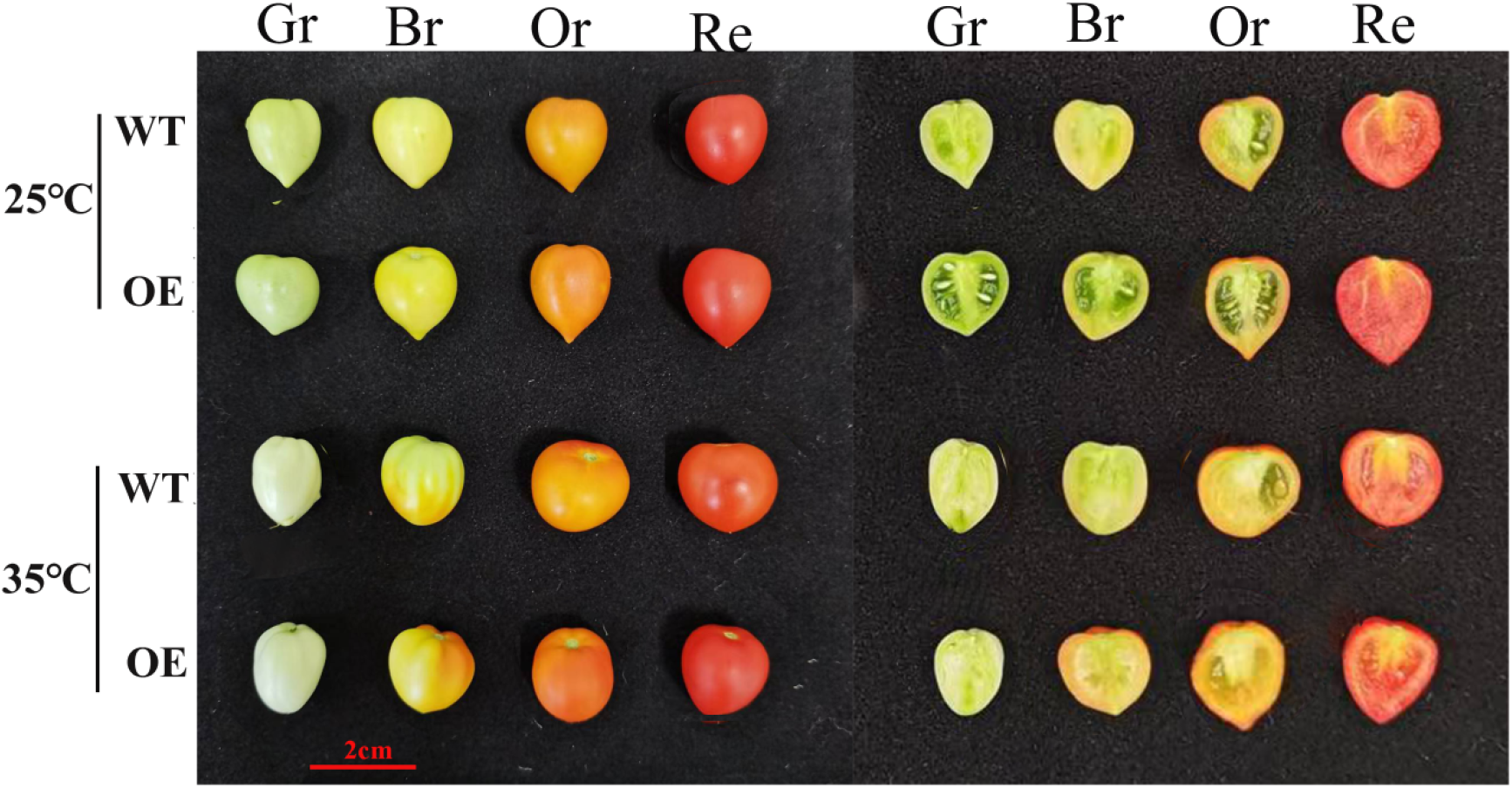
Fruit phenotypic changes in tomato plants under ambient control and high temperature treatments. WT (Wild Type), OE (*SlATG8f -OE)*, Gr(Green ripening),Br(Colour breaking),Or(Orange ripening),Re(Red ripening),scale bar (2 cm).

### Overexpression of *SlATG8f* improves fruit set rate under high-temperature stress

To evaluate the effect of *SlATG8f –*overexpression(*SlATG8f* –OE) on fruit maturation under heat stress, the timing of key developmental stages: Mature green, Breaker, Orange ripe, and Red ripe—was recorded and analyzed (Fig. 2). Under control conditions, all four developmental stages occurred earlier in wild-type (WT) plants than in *SlATG8f* –OE fruits. In contrast, high-temperature stress delayed maturation in WT plants but promoted it in *SlATG8f* –OE fruits, resulting in consistently earlier ripening in the transgenic line compared to WT under stress conditions.

**Fig. 2.**
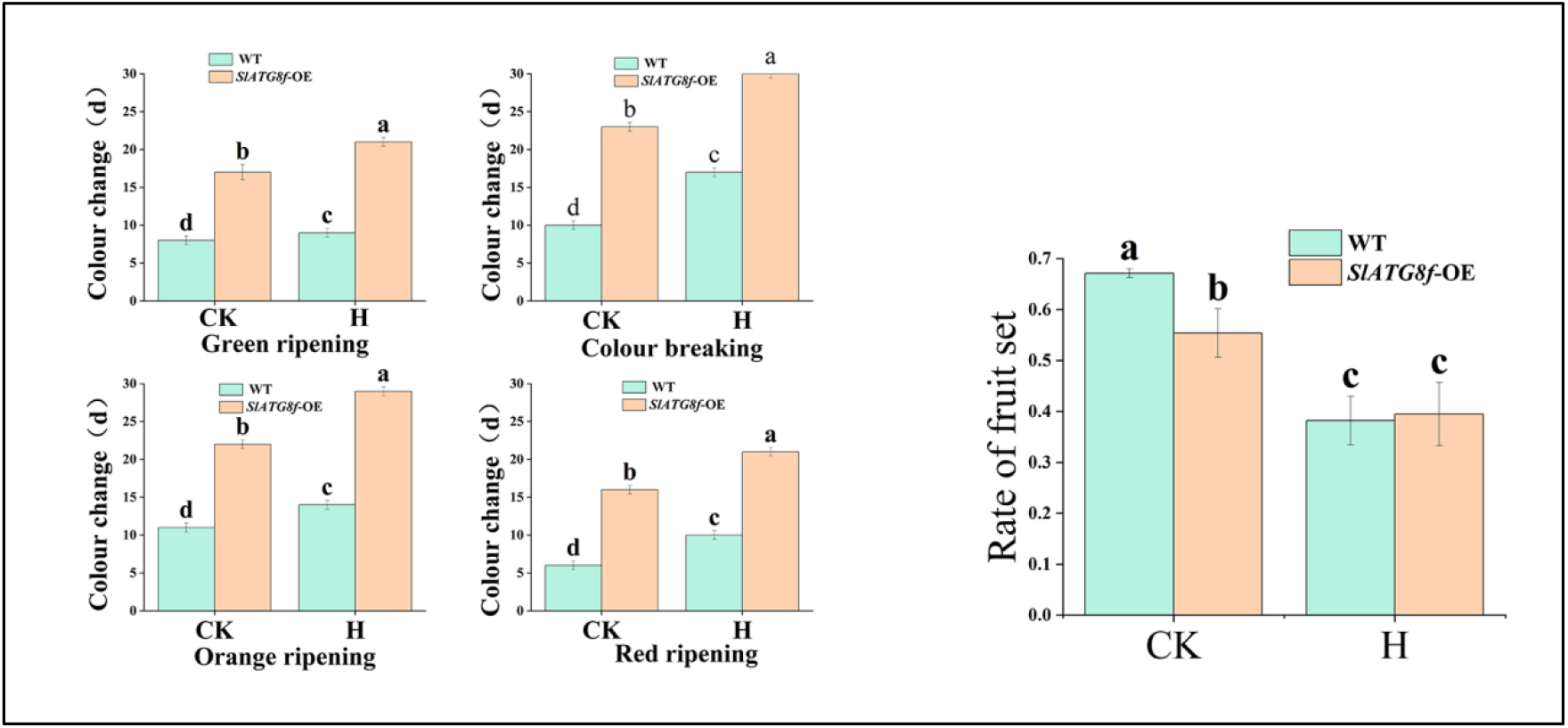
Changes in fruit set in tomato plants under ambient control and high temperature treatments.CK (control), H (high temperature treatment), WT (Wild Type), *SlATG8f -*OE (*SlATG8f* overexpressing plants), (P<0.05)

### Tomato Fruit Quality under Heat Stress Is Modulated by *SlATG8f -OE*

To evaluate the effect of *SlATG8f*-OE on tomato fruit quality under high-temperature stress, key quality parameters were measured in wild-type (WT) and *SlATG8f*-OE fruits under both control and stress conditions (Fig. 3).

**Fig. 3.**
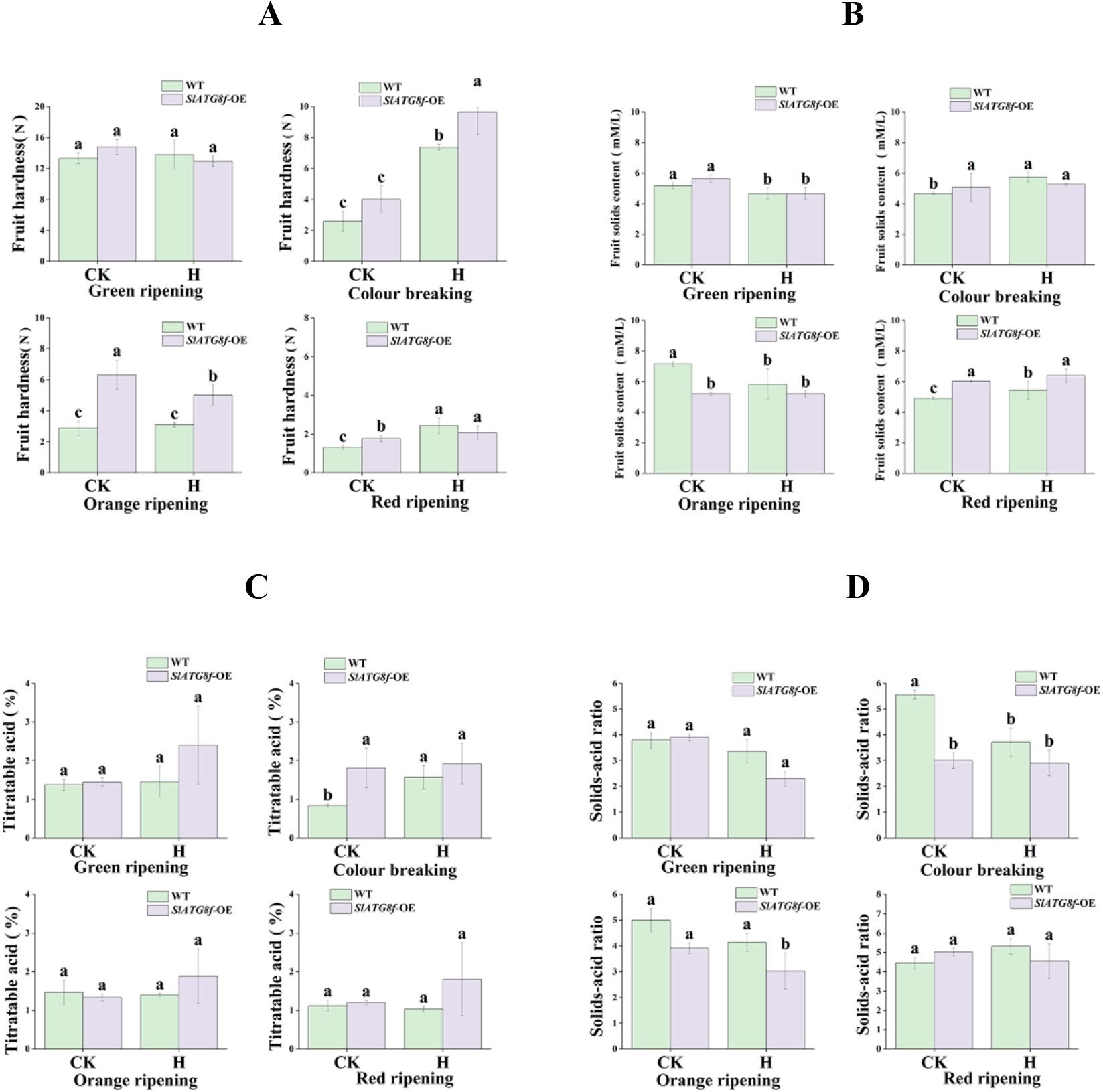
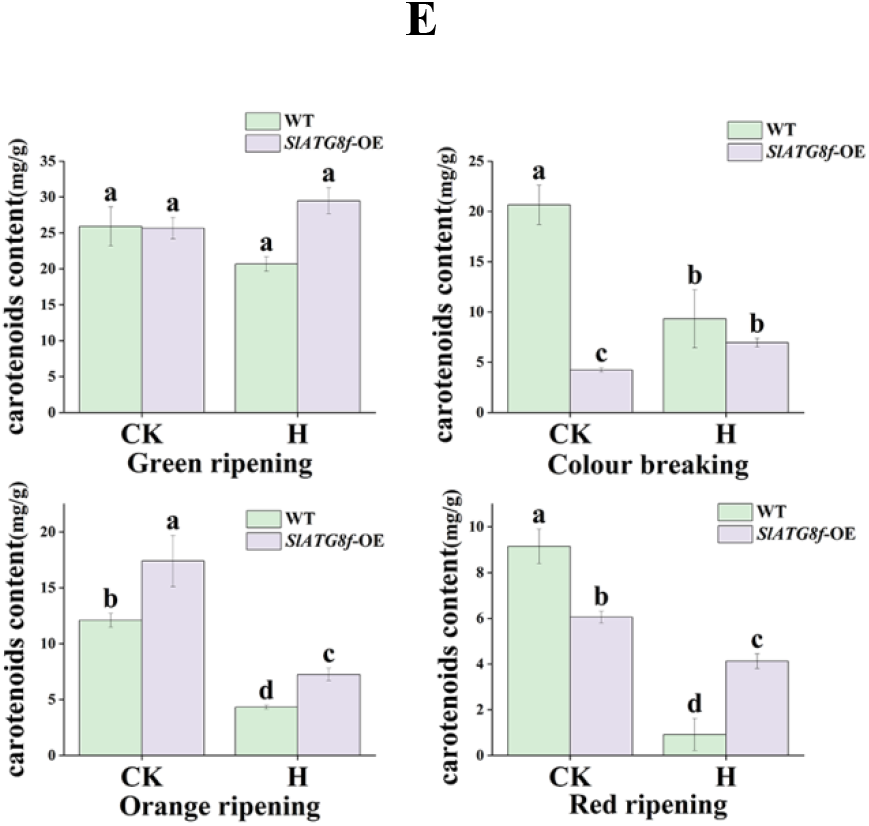
Changes in fruit quality indexes of tomato plants under ambient control and high temperature treatments.CK (control), H (high temperature treatment), WT (Wild Type), *SlATG8f* -OE (*SlATG8f* overexpressing plants), A (fruit hardness), B (fruit soluble solids), C (titratable acid), D (solids-acid ratio), E (carotenoids content), (P<0.05).

Fruit firmness was generally higher in *SlATG8f*-OE fruits than in WT, except at the green ripe stage, where WT fruits were slightly firmer. High temperature further increased firmness in both genotypes (Fig. 3A). Soluble solid content (SSC) was similar between genotypes across treatments; however, heat stress increased SSC at the breaker and red ripe stages while decreasing it at the green and orange ripe stages (Fig. 3B).

Titratable acid (TA) content was consistently higher in *SlATG8f*-OE fruits than in WT under all conditions and was slightly enhanced by high temperature across developmental stages (Fig. 3C). The solid-acid ratio was higher in WT fruits and was reduced under heat stress in both lines (Fig. 3D).

Under control conditions, carotenoid content was higher in WT fruits. However, heat stress reduced carotenoid accumulation in WT at the orange and red ripe stages, resulting in lower levels than in OE fruits under stress (Fig.3E).

Overall, *SlATG8f*-OE fruits exhibited higher firmness, SSC, and TA under high-temperature stress compared to WT.

### *SlATG8f-* Overexpression Modulates Enzyme Activities in Tomato Fruit under High Temperature

To assess the effect of *SlATG8f*-overexpression(*SlATG8f*-OE) on antioxidant enzyme activities under High-temperature stress, key biochemical indicators were measured in fruits from both wild-type (WT) and *SlATG8f****-***overexpressing (*SlATG8f*-OE) plants under control and stress conditions (Fig. 4).

**Fig. 4.**
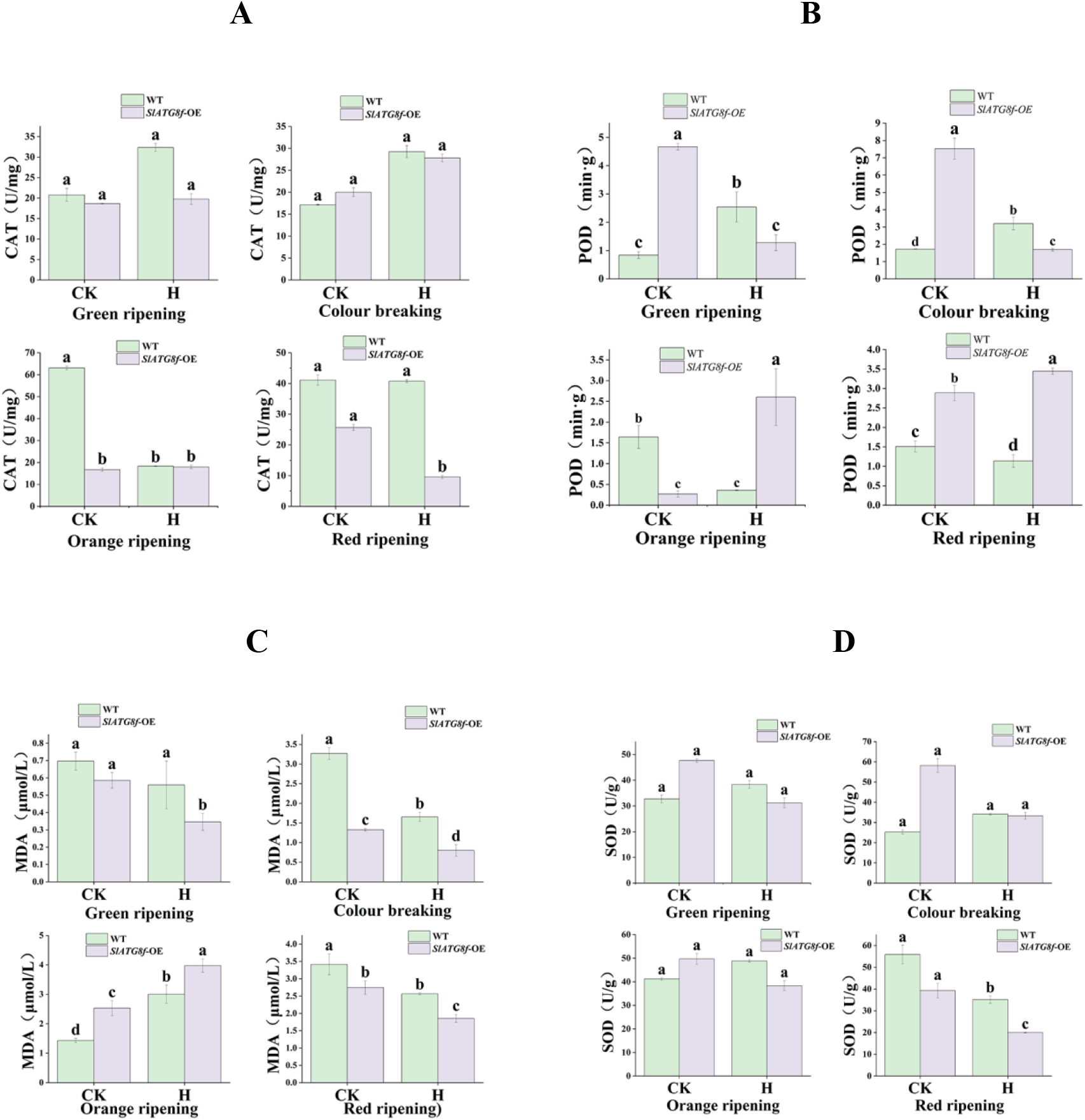
Determination of antioxidant enzyme activities in tomato fruits under ambient control and high temperature treatments.CK (control), H (high temperature treatment), WT (Wild Type), *SlATG8f* OE (*SlATG8f* overexpressing plants), A (CAT enzyme activity), B (POD enzyme activity), C ((MDA content)) D (SOD enzyme activity), (P<0.05)

High-temperature stress increased catalase (CAT) activity, with WT fruits consistently exhibiting higher CAT activity than *SlATG8f*-OE fruits across all developmental stages (Fig. 4A). In contrast, peroxidase (POD) activity was generally elevated in *SlATG8f*-OE fruits compared to WT. Heat stress enhanced POD activity at the orange and red ripe stages but suppressed it at the green and breaker stages (Fig. 4B).

Malondialdehyde (MDA) content was higher in WT fruits than in *SlATG8f*-OE fruits at all stages except orange ripe. Under high temperature, MDA content increased in WT fruits at the orange ripe stage but decreased in the other three stages (Fig. 4C).

Under control conditions, superoxide dismutase (SOD) activity was higher in *SlATG8f*-OE fruits than in WT. However, heat stress enhanced SOD activity in WT fruits while reducing it in *SlATG8f*-OE fruits (Fig. 4D).

### Heat stress modulates the expression of heat-shock protein response genes in Tomato fruit

The expression of heat stress response genes was analyzed in wild-type (WT) and *SlATG8f*-overexpressing (*SlATG8f*-OE) fruits under both control and high-temperature conditions (Fig. 5). Undercontrol conditions, *SlHSFA2* expression was highest in *SlATG8f*-OE fruits at the breaker stage and remained stable under heat stress. In contrast, *SlHSFA3, SlMYB21, and SlMYB26 sh*owed low overall expression, with *SlHSFA3* and *SlMYB26 l*evels higher in WT than in *SlATG8f*-OE fruits. *SlHSFA2* was more highly expressed in WT, whereas *SlMYB21* was elevated in *SlATG8f*-OE fruits. Heat stress upregulated *SlHSP20, SlHSP21*, and *SlHSP70* in both genotypes, with stronger induction observed in WT. Notably, *SlHSP90* expression peaked in WT fruits—at the green ripe stage under control conditions and at the breaker stage under heat stress—and was significantly higher in WT than in *SlATG8f*-OE fruits across all developmental stages.

**Fig. 5.**
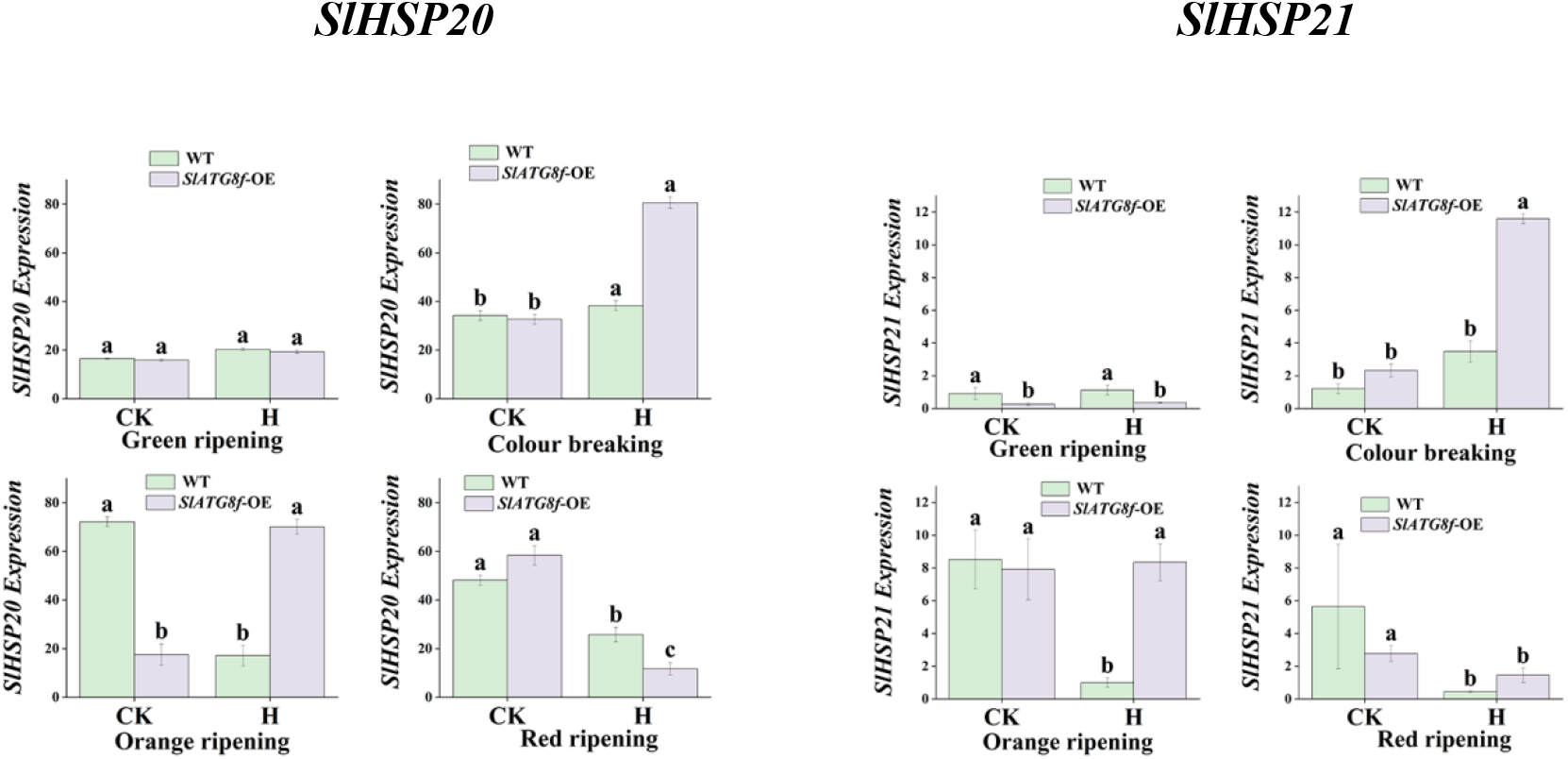
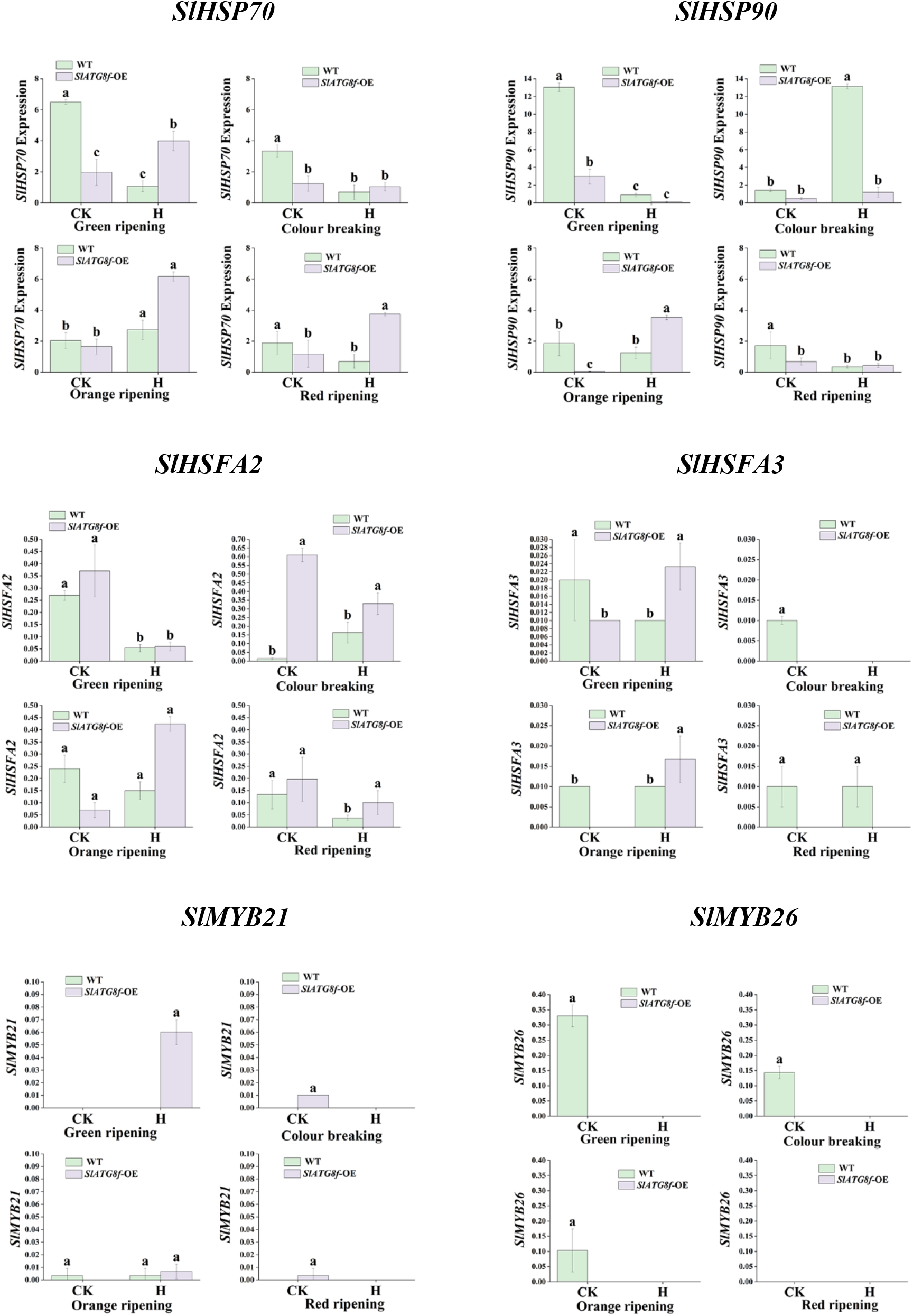
Differential expression analysis of fruit heat stress protein response genes in tomato plants under ambient control and high temperature treatment. CK (control), H (high temperature treatment), WT (Wild Type), *SlATG8f* OE (*SlATG8f* overexpressing plants), (P<0.05).

### *ATG8* Family Gene Expression in Tomato Fruit under Heat Stress

This study aimed to determine whether High-temperature stress enhances autophagic activity in tomato fruit by analyzing the expression of *ATG8* family genes via qRT-PCR (Fig. 6).

**Fig. 6.**
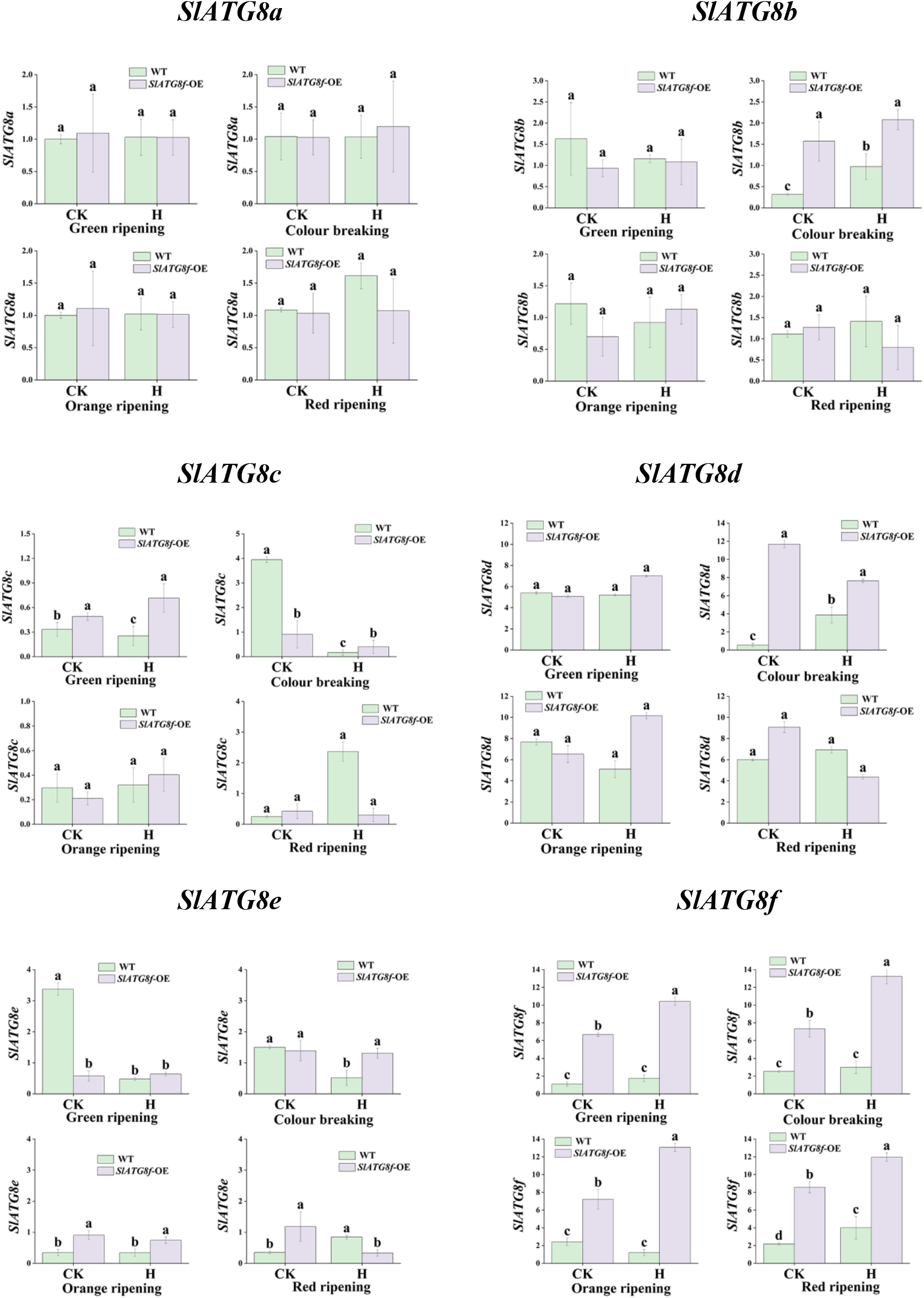
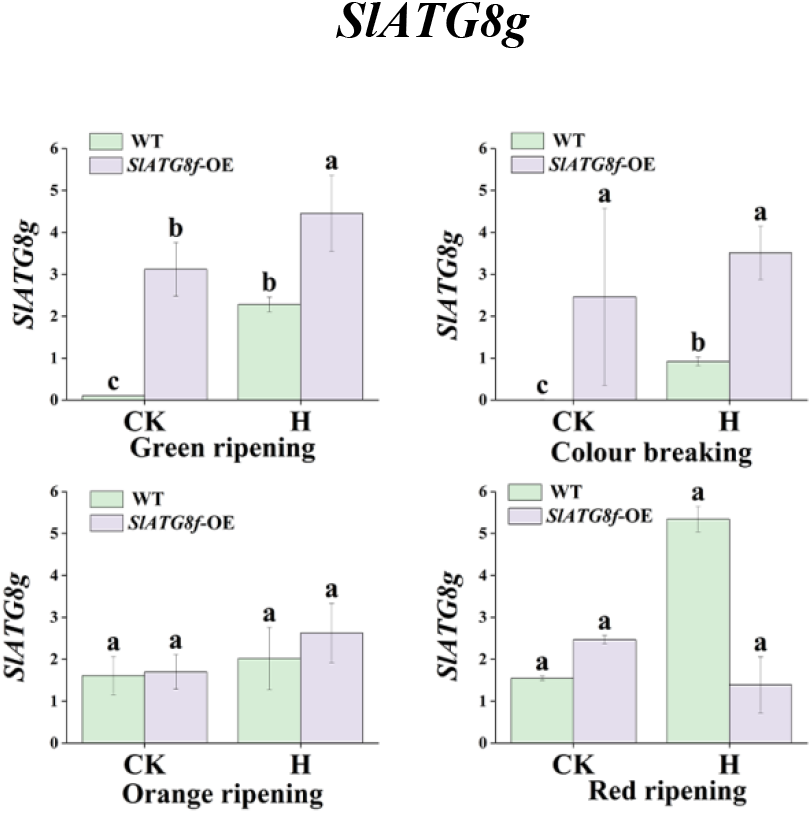
Analysis of differential expression of *SlATG8s* in fruit of tomato plants under ambient control and high temperature treatments.CK (control), H (high temperature treatment), WT (Wild Type), *SlATG8f* OE (*SlATG8f* overexpressing plants), (P<0.05)

The results showed that *SlATG8a* expression remained stable in both WT and *SlATG8f-OE* fruits and was not affected by temperature. *SlATG8b* expression exhibited a unimodal pattern across fruit developmental stages. *SlATG8c* showed its highest expression in WT fruits under control conditions, with relatively stable expression after the breaker stage.

In contrast, *SlATG8d* expression peaked at the breaker stage in *SlATG8f-OE* fruits under both conditions, while remaining consistently low in WT. Overall, *SlATG8d* levels were higher in *SlATG8f-*OE fruits than in WT, following an initial increase and subsequent decrease. *SlATG8e* expression was highest in WT fruits at the green ripe stage under control conditions and remained stable thereafter. *SlATG8f* expression was higher in *SlATG8f-OE* fruits than in WT at the same stage. Meanwhile, *SlATG8g* expression was nearly undetectable at the green and breaker stages in WT control fruits and generally displayed an initial increase followed by a decrease across development.

## Discussion

Fruit development is a crucial phase in plant reproduction, involving coordinated processes such as cell division, expansion, and metabolic changes. In tomato, ripening is characterized by color transition, tissue softening, and the accumulation of various metabolites [21,22], Autophagy, a highly conserved degradation pathway in eukaryotes, has been implicated in fruit metabolism and nutrient recycling during development, although its role remains underexplored [23]. While autophagy has been extensively studied in vegetative tissues across numerous crop species [24], its function in fruits under abiotic stress conditions is still poorly understood. Emerging evidence indicates that autophagy-related genes are upregulated during late ripening stages in grapes and are essential for quality formation in chili and strawberry fruits [25–27], Consistent with these reports, our findings demonstrate that *SlATG8f*-OE promotes locular gel liquefaction and enhances tissue integrity under normal conditions. Under high-temperature stress, however, both wild-type (WT) and *SlATG8f-OE* fruits exhibited morphological deformities and delayed gel development, although the transgenic lines maintained superior firmness, soluble solid content, and titratable acidity. Notably, *SlATG8f-OE* fruits showed higher carotenoid retention under heat stress compared to WT, which suffered a significant reduction—highlighting the protective role of *SlATG8f in* mitigating heat-induced impairment of bioactive compounds. This heat sensitivity in carotenoid accumulation aligns with previous observations in tomato under prolonged high-temperature treatment [28].

Autophagy is recognized as a key response activated under heat stress, facilitating the rem oval of damaged proteins and organelles [29].Mutations in core autophagy genes such as *ATG5* and *ATG7* have been shown to increase thermosensitivity in *Arabidopsis* and tomato [8,30]. highl ighting the importance of autophagy in thermotolerance. The heat stress response involves com plex regulatory mechanisms, including the unfolded protein response and chaperone signaling pathways [29,31]. Our results further demonstrate that overexpression of *SlATG8f* enhances the expression of multiple *ATG8* family members and heat shock proteins (*HSPs*) under high-tempera ture conditions. Although certain HSPs, including *SlHSP90*, exhibited higher expression in wild-type (WT) fruits, most *HSP* and autophagy-related genes were significantly upregulated in *SlAT G8f*-OE fruits. This suggests a coordinated activation of protein quality control pathways media ted by *SlATG8f* under heat stress.

However, there are still some limitations in this study, which also point the way for future in-depth research. Primarily, our conclusions are based on an *SlATG8f* overexpression model. Although overexpression provides valuable phenotypic insights, the generation and analysis of loss-of-function mutants—for instance, using CRISPR/Cas9—would be essential to fully validate *SlATG8f’s* role in thermotolerance. Just as [32] used VIGS technology to demonstrate that *SlATG8f* coordinates ethylene signaling and chloroplast turnover to drive ripening under normal conditions, obtaining reverse genetic evidence—such as through CRISPR/Cas9 mutagenesis—would unequivocally establish the necessity of *SlATG8f* in thermotolerance. In addition, [33]and[34] confirmed the function of autophagy in aging and stress response using *ATG* mutants in Arabidopsis thaliana and tomato, respectively, which provides a reference for us to construct stable mutants in the future.

Secondly, while this study used qRT-PCR data to suggest elevated autophagy activity, autophagy is a dynamic process whose activation is best confirmed through detection of *ATG8* lipidation (*ATG8*-PE)—a hallmark of autophagosome formation. Thus, Western blot analysis of lip id-modified *SlATG8f* protein should be considered the “gold standard” method for verifying enh anced autophagic flux, as strongly recommended in established autophagy monitoring guidelines [35]. In addition,[36]Western blot analysis of *ATG8* lipidation is widely employed in plant autophagy research. Future studies examining dynamic changes in *SlATG8f* protein levels and lipidation would substantially strengthen our conclusions.

Finally, the mechanism by which *SlATG8f* regulates *HSP* gene expression remains unclear. It is unknown whether *SlATG8f* directly interacts with heat shock transcription factors such as *SlHSFA2* or *SlHSFA3*, or whether it enhances the heat shock response indirectly through selective autophagy of negative regulators of the *HSP* pathway. Techniques such as yeast two-hybrid (Y2H), luciferase complementation assay (LCI), or co-immunoprecipitation (Co-IP) could help unravel these interactions. Previous studies offer valuable methodological guidance; for example, the interaction between plant ATG8 and specific target proteins has been well documented[37] and *GmATG8s* were found to regulate leaf aging via interaction with *GmHSP90* in soybean[38] These findings provide a useful reference for investigating *SlATG8f–HSP* interactions in tomato.

In conclusion, this study demonstrates that *SlATG8f* positively contributes to thermotolerance in tomato fruit under high-temperature stress, providing a valuable genetic resource and theoretical foundation for improving crop stress resistance and fruit quality through genetic engineering. Future research should focus on generating stable loss-of-function mutants, direct measurement of autophagic flux at the protein level, and elucidating the interaction between *SlATG8f* and heat shock signaling components to fully unravel the molecular network mediated by this gene

## Conclusions

In this study, wild-type (WT) and *SlATG8f*-overexpressing (*SlATG8f* -OE) tomato plants were subjected to high-temperature stress (35 °C), with 25 °C serving as the control. Comparative analyses revealed that *SlATG8f*-OE enhanced thermotolerance during fruit development, as reflected by improved phenotypic traits, accelerated fruit set and color transition, and promoted maturation. Furthermore, *SlATG8f*-OE elevated the expression of several *ATG8* family members and heat shock protein-related genes under high-temperature conditions.

These results indicate that *SlATG8f* overexpression enhances autophagic activity and activates heat stress response pathways, thereby promoting early fruit ripening and improving fruit quality under thermal stress. Our findings underscore the critical role of *SlATG8f* ***i***n regulating tomato fruit development under abiotic stress and provide a theoretical foundation for improving tomato quality and stress resilience in breeding programs.

## Declaration of Competing Interest

The authors declare that they have no known competing financial interests or personal relationships that could have influenced the work reported in this paper.

## Acknowledgments

The authors are grateful to the National and Local Vegetable Engineering Center (Guizhou) and the Laboratory of the Department of Horticulture (Agricultural College of Guizhou University) for their support to this project.

## Author contribution statement

Qunmei Cheng: Conceptualization, Writing original draft, Formal analysis, Data curation, Software, Methodology, Validation. Cen Wen: Software, Validation. Zhuo He : Software, Validation. Liu Song: Software, Validation. Wen Xu: Writing - review & editing, Conceptualization, Funding acquisition, Project administration, Resources, Supervision. All authors have read and agreed to the published version of the manuscript.

## Data availability

No data was used for the research described in the article.

## Funding

This work was supported by the National Natural Science Foundation of China (31760594 and 32260754), the Special Project for Cultivating Innovative Talents of Thousand Levels in Guizhou Province[(2018)02], the Guizhou University of Science and Technology Research Start-up Fund for New Faculty [(2016)48].

## Appendix A. Supplementary data

Supplementary Table S1 name primers for each gene and Internal reference gene Actin (Solyc03g078400), Table S2 name qRT-PCR reaction system and reaction program.

## Notes

### Competing Interest Statement

The author declares no conflict of interest

